# Local translational repression and retention of SynGAP1 during synaptic plasticity

**DOI:** 10.64898/2026.04.29.719691

**Authors:** Rhys W. Livingstone, Paul G. Donlin-Asp

## Abstract

Synaptic plasticity depends on tightly controlled protein production, yet most locally synthesised synaptic proteins studied to date promote strengthening. Here, we investigate whether SynGAP1—a key negative regulator of Ras/Rap signalling—undergoes local translation to constrain plasticity. We find highly regulated developmental distribution of SynGAP1 isoforms, with stable synaptic occupancy of α1, progressive enrichment of α2, and late dendritic accumulation of β, with all isoforms positioned at the PSD. Despite abundant dendritic *Syngap1* mRNA, nascent SynGAP1 translation is dynamically regulated, showing suppression during synaptic maturation. Contrary to the prevailing model, we find that activity-induced dispersion of SynGAP1 is an artefact of GFP overexpression, as the endogenous protein remains stably anchored at synapses following LTP induction. Instead, we uncover a transient, dendrite-specific suppression of SynGAP1 synthesis upon LTP induction. Our findings oppose the current model of SynGAP1 dispersion and identify translational repression as a key regulatory mechanism for synaptic SynGAP1. We propose that synapses transiently lift inhibitory constraints not by physically removing SynGAP1, but through its well reported inactivation and halting its local production.

## Introduction

Learning and memory are fundamental processes that allow the brain to acquire, retain, and retrieve information. Central to the storage of information in the brain are its trillions of synapses – the physical connections between neurons where information in the brain is stored. Molecular and cellular changes within the brain contribute to specific modifications in the strength and efficacy of synapses across development and in response to stimulation, through synaptic plasticity. One such process is that of long-term potentiation (LTP), the activity-dependent enhancement of synaptic efficacy. One of the key mechanisms involved in the manifestation and maintenance of LTP, and in the consolidation of memories, is protein synthesis^1,2,3^, which is believed to supply the cohort of proteins required during LTP. Crucially, synapses can be hundreds of microns from the cell body, making their local regulation and protein supply a critical logistical challenge for the neuron. To overcome this, neurons localise thousands of mRNAs to synapses and engage in local protein production on demand, allowing localised adaptability^4^. Numerous forms of synaptic plasticity utilise the recruitment of mRNA to synapses^5,6^, and undergo local protein synthesis for their manifestation or maintenance^1,7–10^. So far, typically studied locally synthesised proteins tend to be those which exert a positive function in plasticity, such as those which advantage the strengthening of the postsynaptic response; postsynaptic density protein 95 (PSD95)^5^, Ca^2+^/calmodulin-dependent protein kinase II (CaMKII)^11,12^, the α-amino-3-hydroxy-5-methyl-4-isoxazolepropionic acid (AMPA) receptor subunit GluA1^13^, β-actin^6^, and small GTPases including Rac1^14^. However, synaptic plasticity involves both strengthening and weakening connections, so it remains unlikely that only proteins involved in strengthening synapses would be produced. Examples exist of locally synthesised proteins whose functional role involve the negative regulation of processes such as growth cone growth and guidance; RhoA, a GTPase locally synthesised in the growth cone of developing axons negatively regulates microtubule protrusion^15,16^, indicating a similar role may exist for negative regulators in synaptic plasticity.

One candidate for a locally synthesised negative regulator is SynGAP1 (Synaptic GTPase-Activating Protein 1), a Ras/Rap GAP enriched at excitatory neuronal synapses^17^, of which *Syngap1* mRNA has been found to be present in the rat hippocampal neuropil (Glock et al. 2021), and neurites of cultured iNeurons^18^. Encoded by the *SYNGAP1* gene, *Syngap1* mRNA exists as three primary N-terminal isoforms (SynGAP1 A, B, and C), each of which can be alternatively spliced into a further four isoforms, differing in their C-termini; α1, α2, β, and γ, each proposed to possess unique functions. The presence of SynGAP1 at synapses is thought to act to negatively regulate the manifestation of synaptic plasticity during development^19^, through tonic inhibition of Ras/Rap activity^20,21^, and, due to the presence of the PDZ binding domain in SynGAP1α1^22^, occupies ‘slots’ within the postsynaptic density (PSD), effectively preventing AMPA receptor (AMPAR) insertion and spine growth^23,24^. The importance of this regulation has been highlighted through SynGAP1 mutations, leading to a functional haploinsufficiency of SynGAP1, and resulting in enlarged spines^25^, impaired LTP^19^, and elevated Ras/ERK signalling^26^, as well as higher-order changes such as excitatory and inhibitory (E/I) imbalances^27^, and altered cortical circuit assembly^28^. In humans, these impairments manifest as SYNGAP1-related intellectual disability, a disorder characterised by neurodevelopmental issues, including autism spectrum disorder, intellectual disability, and epilepsy^29^.

At the synapse, and upon N-methyl-D-aspartate (NMDA) receptor activation, phosphorylation of SynGAP1 by CaMKII alleviates the inhibition of Ras/Rap^21^, permitting the induction of LTP and downstream processes. This process has previously been described as involving the physical dispersion of SynGAP1 protein from the PSD^30^ and the spine^20^. Importantly, following induction of LTP, potentiated synapses enter a refractory period during which further strengthening is impeded^31,32,33^. During this, processes driving postsynaptic strengthening, such as synaptic AMPAR accumulation and spine growth, approach a functional ceiling, during which SynGAP1 protein is thought to return to the synapse, and halt further synaptic potentiation. However, it is currently unclear how and when SynGAP1 return is mediated. With the presence of *Syngap1* mRNA at distal sites, the return of SynGAP1 to the synapse could be driven by the local synthesis of *de novo* SynGAP1 protein during LTP. Here, we explore the hypothesis that the local synthesis of SynGAP1 restores synaptic plasticity by supplying a pool of *de novo* SynGAP1 protein at the synapse.

## Results

### SynGAP1 isoforms α1, α2, and β protein accumulate in the dendrite and at the synapse throughout neuronal development *in vitro*

Due to the strong relationship between typical neurodevelopment and the regulation and expression of SynGAP1, we first sought to assess the dendritic and synaptic accumulation of endogenous SynGAP1. We focused on three of the C-terminal SynGAP1 isoforms at key developmental stages in cultured neurons at 7, 14, and 21 days in vitro (DIV), a period spanning neuronal and synaptic development and maturation^34^. To evaluate this quantitatively, we analysed the number and intensity of individual SynGAP1 puncta throughout the dendrites, relative to the excitatory synaptic protein PSD95. Across development, PSD95 accumulation significantly increased from DIV7 through to DIV14 and DIV21 (Fig. 1 A-C), indicating an incremental increase in the number of new synapses. Conversely, the synaptic accumulation of PSD95 protein remained stable between DIV7 and DIV14, increasing at DIV21, indicating a late accumulation of PSD95 protein at mature synapses. Both the dendritic and synaptic accumulations of SynGAP1α1 remained constant across all developmental stages (Fig. 1 A,D & E), likely indicating a highly conserved synaptic coverage and copy number for SynGAP1α1 at both young and mature synapses. Comparable to PSD95, the number of SynGAP1α2-positive synapses increased across development, peaking in mature neurons, while the synaptic accumulation remained constant across development (Fig. 1 A,F & G). In contrast, SynGAP1β abundance remained stable between DIV7 and DIV14, with dendritic accumulation increasing exclusively by DIV21, possibly indicating a role in the development or stabilisation of mature synapses. Synaptic SynGAP1β protein accumulation also remained stable across development, indicating that SynGAP1β levels may be maintained at consistent levels throughout development (Fig. 1 A, H & I). While mature neurons showed a slightly greater number of SynGAP1β puncta relative to SynGAP1α1 and α2 isoforms (Fig. 1 D, F & H), α2 showed consistently greater synaptic accumulation across dendritic development, suggesting that SynGAP1α2 may be the dominant isoform at the synapse. The greater number of SynGAP1β puncta likely reflects non-synaptic accumulation, in line with previous reports showing higher cytosolic SynGAP1β than other isoforms^22^.

**Figure 1.**
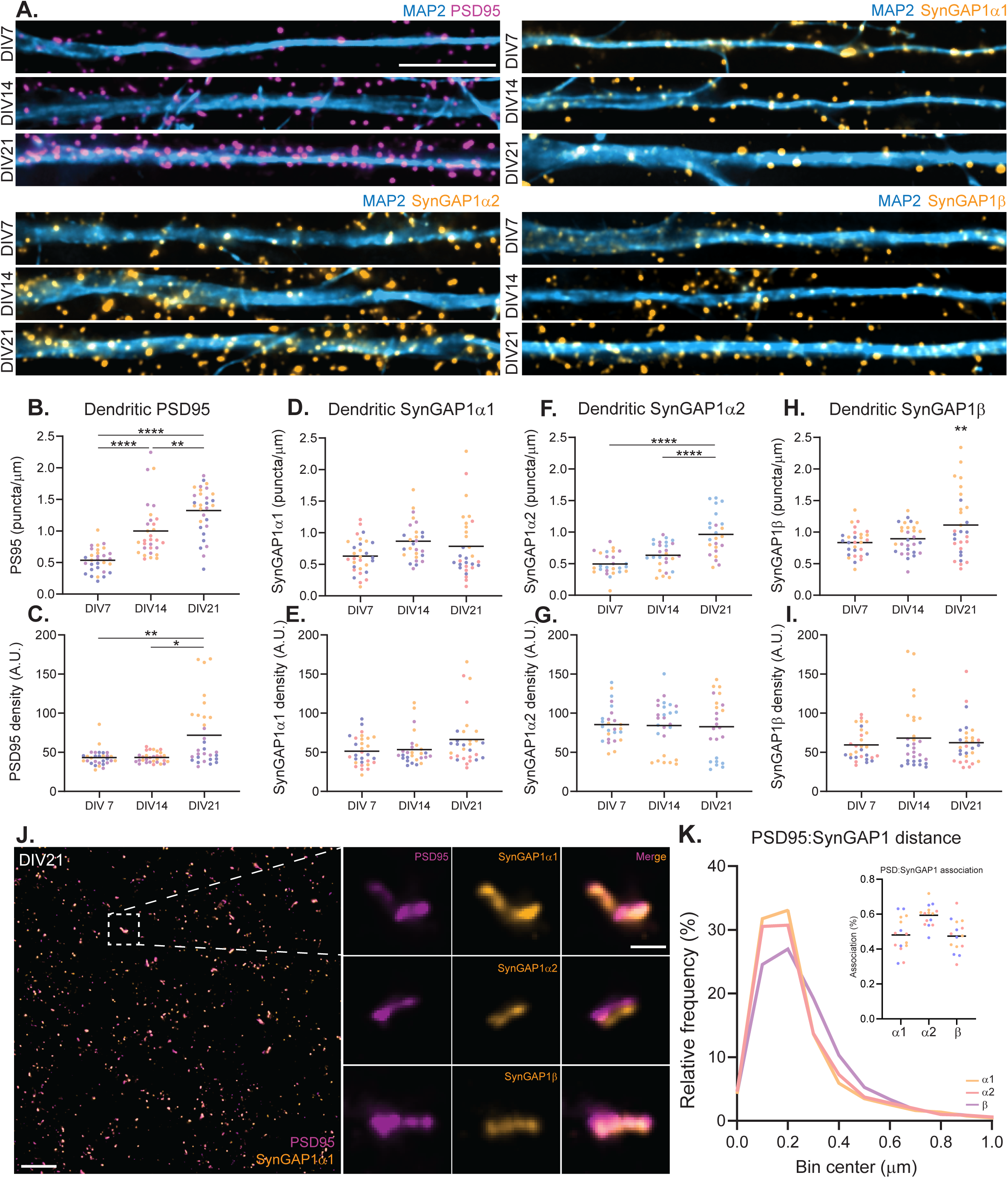
SynGAP1 isoforms accumulate at the synapse. **A)** Representative images show accumulation of PSD95 protein (top left panels, magenta), SynGAP1α1 (top right, orange), SynGAP1α2 (bottom left, orange), and SynGAP1β (bottom right, orange) in developing dendrites (MAP2, blue). Quantification of **B)** PSD95 puncta per µm and **C)** puncta intensity (*n* = 3 biological replicates, 30 dendrites). **D)** SynGAP1α1 puncta per µm E) and puncta intensity remains relatively stable across DIV7, 14, and 21 (*n* = 3 biological replicates, 27-30 dendrites). SynGAP1α2 increases in the number of **F)** dendritic puncta, **G)** puncta density remains stable (*n* = 3 biological replicates, 24-26 dendrites). The number of SynGAP1β-positive **H)** dendritic puncta peaks at DIV21 while **I)** puncta density remains stable throughout (*n* = 3 biological replicates, 29 dendrites). Significance was determined by Ordinary one-way ANOVA or Kruskal–Wallis one-way ANOVA with Dunn’s multiple comparisons test where appropriate. Normality for all data was determined by Shapiro-Wilk normality tests. *p* ≤ 0.05*, *p* <0.01**, *p* <0.0001****. Scale bar = 10 µm. **J)** Ten-fold expansion microscopy reveals the distribution of SynGAP1α1 (orange) relative to PSD95 (magenta), scale bar = 10 µm; magnified inset shows a single synapse containing SynGAP1α1, middle and lower panels show single synapse examples of SynGAP1α1 and SynGAP1β, respectively. Scale bar = 1 µm. **K)** Quantification of SynGAP1α1 (orange, *n* = 3 biological replicates, 5695 synapses), SynGAP1α2 (pink, *n* = 3516 synapses), and SynGAP1β (purple, *n* = 1902 synapses) isoform distributions relative to PSD95, inset shows relative association of SynGAP1 isoforms with PSD95. All data points are coloured to represent data from 3 independent biological replicates.

### Synaptic localization of SynGAP1 isoforms

SynGAP1α1 is the only isoform to contain a PDZ-binding domain, yet we have shown here, and in line with previous observations^22^ α2 and β isoforms show discrete punctate localisation, consistent with synaptic localisation throughout the dendrites of cultured hippocampal neurons. To further examine the relationship between SynGAP1 isoforms and their proximity to PSD95, we employed super-resolution ten-fold expansion microscopy^35^ (Fig. 1). All isoforms are positioned in close proximity to PSD95 (Fig. 1 J-K; SynGAP1α1: 26.6 nm ± 0.29 centroid-centroid (2.66nm expansion corrected); SynGAP1α2: 24.8 ± 0.28 (2.48nm expansion corrected); SynGAP1β: 28.0 ± 0.22 (2.80nm expansion corrected), these indicate a very uniform localisation pattern of SynGAP1 isoforms at the synapse. On average, the proportion of SynGAP1 associated with PSD95 was 48.09% ± 0.10, 59% ± 0.061, and 47.4% ± 0.093 for SynGAP1α1, SynGAP1α2, and SynGAP1β, respectively, likely indicating an overlapping occupation of these isoforms across the synaptic population. Consistent with higher protein expression (Fig. 1 A, F & G) for SynGAP1α2 relative to the other isoforms, α2 shows the highest association with PSD95 (Fig. 1 K), and likely represents the dominant isoform in this neuronal population.

### Nascent SynGAP1α2 protein is dynamically regulated despite unchanging mRNA levels

Next we sought to assess whether the developmental accumulation of SynGAP1 is driven by local protein synthesis, assessing both mRNA and nascent protein levels in both the soma and dendrites of developing neurons. In line with previous research^4,18^ using RNA-FISH, we detected both *Syngap1* and *Psd95* mRNA transcripts in developing neurons in culture (Fig. 2 A-E.). In both the soma (Fig. 2 A-C) and dendrites (Fig. 2 A, D-E), *Syngap1* and *Psd95* transcripts showed an increasing developmental trend in transcript abundance, indicating consistent expression of both transcripts despite increased protein accumulation in the dendrites (Fig. 1 A-I).

**Figure 2.**
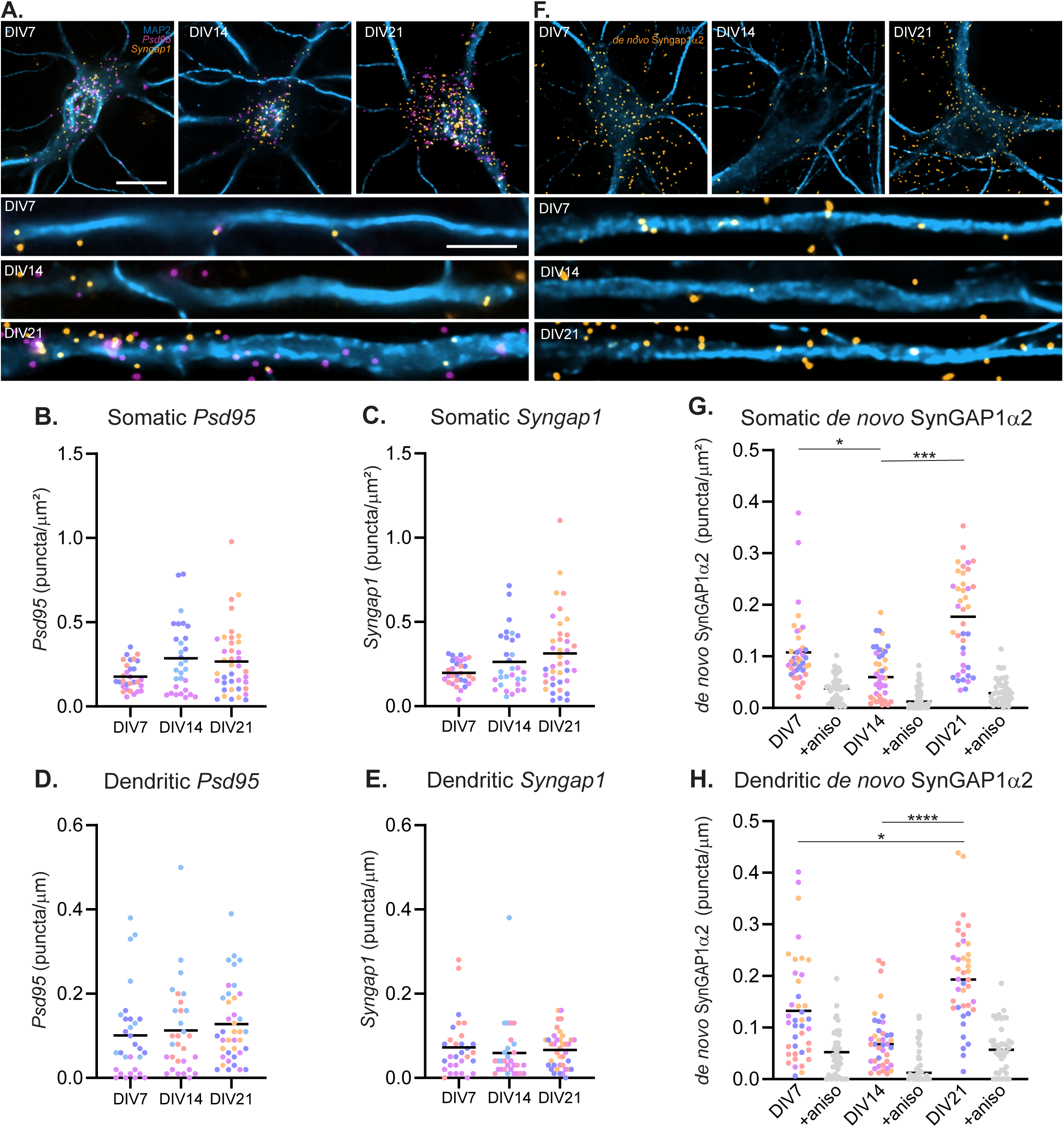
SynGAP1 mRNA and nascent protein throughout development. Representative images and quantification show **A)** steady-state expression of *Psd95* (magenta) and *Syngap1* (orange) mRNA in developing soma (top panels) and dendrites (bottom panels) from DIV7, 14, and 21 neurons (MAP2, blue). *n* = 30-39 soma/dendrites. Quantification shows no change in somatic or dendritic *Psd95* **(B & D)** or *Syngap1* **(C & E)** transcripts. Significance was determined by Kruskal–Wallis one-way ANOVA. **F)** Representative images of nascent SynGAP1α2 (orange) protein in developing soma (top panels) and dendrites (bottom panels; MAP2, blue). Scale bars = 10 µm. Quantification of **G)** somatic and **H)** dendritic nascent SynGAP1α2. *n* = 40-41 soma/dendrites. Normality for all data was determined by Shapiro-Wilk normality tests. Significance was determined by Ordinary one-way ANOVA. *p* ≤ 0.05*, *p* <0.001***, *p* <0.0001****. All data points are coloured to represent data from 3-4 independent biological replicates.

As we find abundant *Syngap1* mRNA across dendritic development, we next examined the developmental pattern of expression for nascent SynGAP1α2, the dominant isoform of SynGAP1, in both the somatic and local dendritic population (Fig. 2 F-H). Unexpectedly, compared to DIV7, somatic nascent SynGAP1α2 was significantly decreased at DIV14, returning to DIV7 levels by DIV21 (Fig. 2 F & G). This trend was further observed in the dendrites between DIV7 and DIV14, while dendritic nascent SynGAP1α2 was found to increase at DIV21, beyond that of DIV7 and DIV14 levels (Fig. 2 F & H). These results indicate a developmental period during which SynGAP1α2 is translationally repressed, likely arising through increased translational regulation. This translational suppression aligns with a period of increased spontaneous activity observed in neuronal cell culture^36,37^, suggesting a relationship between increased synaptic activity and translational suppression of SynGAP1α2.

### GFP-SynGAP1α1 exhibits an all-or-none pattern of dispersion

Both SynGAP1α1 and SynGAP1α2 have been previously reported to disperse from the synapse following cLTP-induced neuronal stimulation^20,22^, and are the principle data supporting a model of physical removal of SynGAP1 from the synapse as a permissive mechanism driving synaptic plasticity. Therefore, to address our primary hypothesis that local *de novo* SynGAP1 synthesis drives the return of SynGAP1 to the synapse, we assessed the timeline of SynGAP1 dispersion, focusing our attention on SynGAP1α1, the isoform thought to have the greatest tendency towards dispersion^20,22^ (Fig 3). Following the induction of cLTP (Fig. 3 A-B & D), we observed an average reduction in puncta signal of ∼25% relative to baseline, which did not return within 60 minutes and instead continued to decline by ∼7% during the post-induction recovery phase.

**Figure 3.**
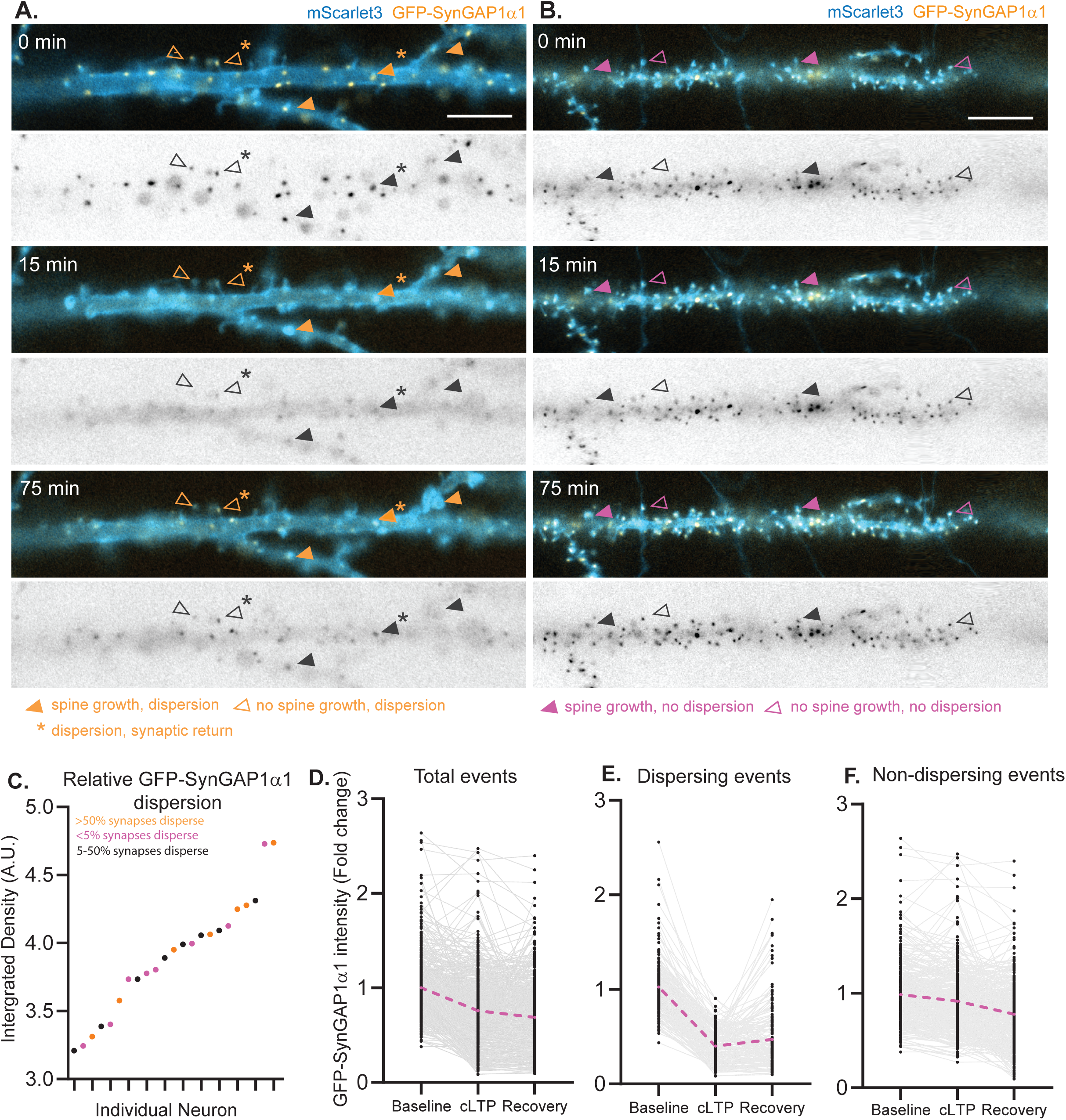
Life cell imaging of GFP-SynGAP1α1 shows an all-or-none pattern of dispersion. Representative time series images of **A)** dispersing and **B)** non-dispersing GFP-SynGAP1α1 (orange) in mScarlet3-expressing neurons (blue) at baseline (0 min, top panels), after cLTP induction (15 min, centre panels), and end of the recovery period (75 min, bottom panels). Orange arrows indicate example spines at which dispersion occurred in the presence (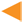) or absence (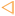) of notable spine size changes. Asterisks indicate spines at which the return of GFP-SynGAP1α1 during the recovery period was observed. Magenta arrows indicate example spines at which dispersion did not occur in the presence (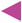) or absence (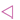) of notable spine size changes. Black & white panels show raw GFP-SynGAP1α1 signal **C)** Individual neurons categorized by the proportion of synapses dispersing relative to the level of GFP-SynGAP1α1 expression at baseline. Data were log transformed for ease of visualisation. **D)** Quantification of dispersion events from all recordings (*n* = 23 neurons, 571 synapses), average is indicated by a dashed magenta line. **E)** Quantification of dispersion events as defined as a minimum of 40% decrease in GFP-SynGAP1α1 signal during the induction of cLTP (*n* = 176 synapses). **F)** Quantification of non-dispersing events defined as a decrease of less than 40% GFP-SynGAP1α1 signal during the induction of cLTP (*n* = 396 synapses). All data is represented as a fold change from control.

To further parse the proportion of synapses at which dispersion occurred, we defined dispersion as a minimum 40% reduction in baseline signal, based on previous observations^20,22^. Crucially, in doing so, we uncovered three primary populations of GFP-SynGAP1α1 responses (Fig. 3 C, E-F). During these experiments, a subset of cells (7/23) showed that more than 50% of synapses dispersed GFP-SynGAP1α1 (Fig. 3 A, C & E, Supplemental video 1; 123/149 synapses), with 2/7 of these exhibiting complete, cell-wide dispersion (53/53 synapses). A further 8/23 cells showed little to no dispersal across all puncta analysed (Fig. 3 B, C & F, Supplemental video 2), showing less than 5% of synapses dispersing GFP-SynGAP1α1, comprising 6/8 cells which showed no detectable dispersion. Lastly, 8/23 cells dispersed greater than 5% but less than 50% (Fig. 3 C-F; comprising 54 of 412 synapses analysed), which may fall more in line with previous observations^20,22^. As expected, synapses that did not disperse by more than 40% of baseline signal during cLTP induction showed, on average, a ∼9% reduction in signal, which continued to decrease during post-cLTP recovery, likely due to photobleaching of GFP. Following dispersion from the synapse, previous accounts describe most synapses as reliably dispersing GFP-SynGAP1α1 for up to an hour after cLTP induction^20,22^. Here, we observed that, in synapses with greater than 40% dispersion of baseline signal, 21.1% (37/177) of synapses returned to within 30% of baseline fluorescence (Fig. 3 A & E), indicating an early return of SynGAP1α1 at these synapses. To confirm cLTP was robustly induced in our cultures, we verified AMPA receptor exocytosis using SEP-GluA1 (Supplemental Fig. 2, Supplemental video 3)^38^.

Notably, there was an observed increase in the propensity of GFP-SynGAP1α1-expressing cells towards dendritic blebbing following cLTP induction (Supplemental Fig. 3 A-C, Supplemental video 4; 30.3%, 10/33 neurons), indicative of excitotoxic cell stress^39^, compared to neurons expressing the control construct SEP-GluA1 (13.3%, 8/60; Supplemental Fig. 3 C). Overexpression of GFP-SynGAP1α1 showed no strong relationship in the propensity of neurons to show dispersion (Fig 3. C), nor cell stress (Supplemental Fig. 3 B), meaning the determining factor for the degree of SynGAP1α1 dispersion remains unknown. However, as SynGAP1 is a crucial hub for downstream signalling cascades, SynGAP1α1 overexpression should be considered for both the manifestation of synaptic plasticity and the maintenance of neuronal health within a physiological range. Overall, these results describe a nuance to SynGAP1α1 dispersion not previously reported^20,22^, and is most consistent with GFP-SynGAP1α1 dispersion resulting from both an artefact of GFP tagging and overexpression.

### Endogenous SynGAP1 does not disperse from the synapse following induction of cLTP

As we and others have shown that SynGAP1 isoforms exhibit strikingly high synaptic abundance^40^, it is feasible that this co-occupancy results in their competition for PSD95 binding, through direct or indirect interactions, and regulation of Ras activity. Due to this, it is likely that overexpression of a GFP-tagged form of SynGAP1 protein may alter the expression of synaptic plasticity, by interfering with endogenous SynGAP1 isoform occupancy at the synapse and levels far higher than normal. GFP-SynGAP1 dispersion, therefore, may be a consequence of overexpression, rather than an actual biological phenomenon. Accordingly we aimed to examine if it is possible to detect dispersion of endogenous SynGAP1 protein (Fig. 4). Following cLTP induction, we probed hippocampal cultures for SynGAP1 α1, α2, and β isoforms and found no change compared to control in either the intensity (Fig. 4 A & B,D,F) or the number of puncta (Fig. 4 A & C,E,G) of SynGAPα1, α2, or β isoforms. These data are inconsistent with SynGAP1 dispersion and are most consistent with synaptic retention of SynGAP1 at the synapse. However, we cannot exclude a more subtle relocalization of SynGAP1 within the synapse, as previously reported with depolarization and immunogold labeling^30^.

**Figure 4.**
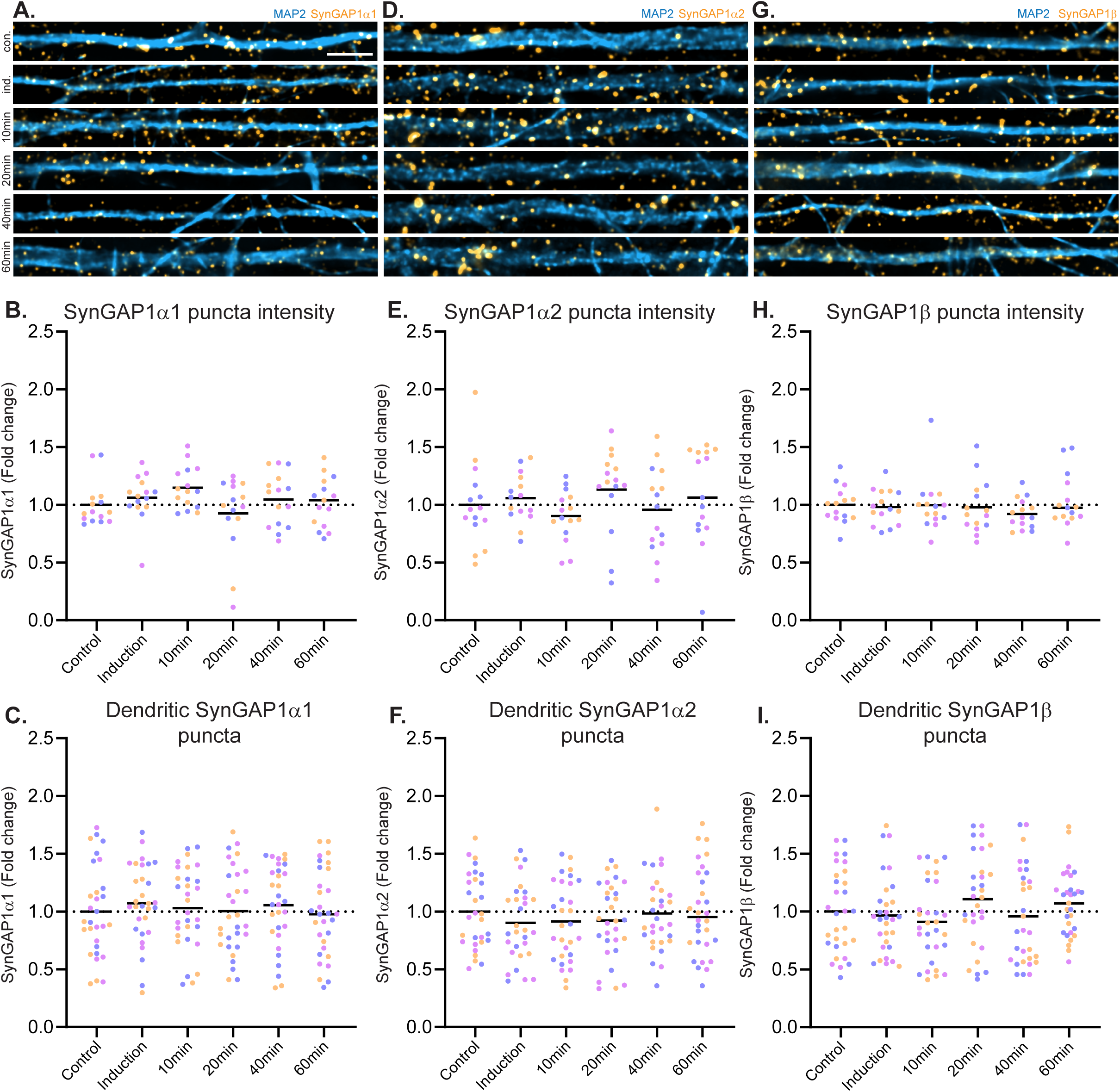
Endogenous SynGAP1 does not disperse at the dendritic level. Representative images show **A)** expression of SynGAP1α1 (left panels, orange), SynGAP1α2 (centre panels, orange), and SynGAPβ (right panels, orange) following the induction of cLTP and recovery across 10-, 20-, 40-, and 60 minutes in neuronal dendrites (MAP2, blue). Quantification showed no changes in **B)** SynGAP1α1, **D)** SynGAP1α2, and **F)** SynGAPβ puncta intensity when analysing the dendritic field of view (FOV, *n* = 15 FOV, 3 biological replicates). Nor were any changes detected in the number of **C)** SynGAP1α1, **E)** SynGAP1α2, and **G)** SynGAPβ puncta per micron in the dendrites of cultured neurons, compared to controls. *n* = 30 dendrites, 3 biological replicates. Scale bar = 10 µm. Significance was determined by Ordinary one-way ANOVA or Kruskal–Wallis one-way ANOVA with Dunn’s multiple comparisons test where appropriate. Normality for all data was determined by Shapiro-Wilk normality tests. All data points are coloured to represent data from 3-4 independent biological replicates.

To verify synaptic retention of SynGAP1, we assessed if cLTP induction has any detectable effect on the subcellular localisation of SynGAP1 within synapses or the number of synapses associated with SynGAP1, through ten-fold expansion microscopy. Here, we found no change in the association of PSD95 with SynGAP1α1 (Fig. 5 A-B), α2 (Fig. 5 C-D), or β (Fig. 5 E-F) isoforms at the synapse. Nor were there any detectable changes in the distance of each isoform from PSD95 (Fig. 5 B, D & F). These results conclusively demonstrate SynGAP1 is retained at the synapse following induction of LTP, nor show any evidence for more subtle relocalizations within the synapse. To validate these findings biochemically, we isolated synaptosomes 10 minutes after cLTP induction. Using this method, we were able to detect a modest, but nonsignificant decrease in synaptic SynGAP1α1 protein relative to PSD95 (Fig. 5 G-H). These data indicate that there may be a subtle change in the affinity of SynGAP1α1 for PSD95 following the induction of cLTP, which does not manifest unless the synapse is functionally isolated from the rest of the neuron. Altogether, these results indicate that SynGAP1 dispersion is a phenotype exclusive to the GFP-tagged protein, and not reflective of the behaviour of endogenous SynGAP1.

**Figure 5.**
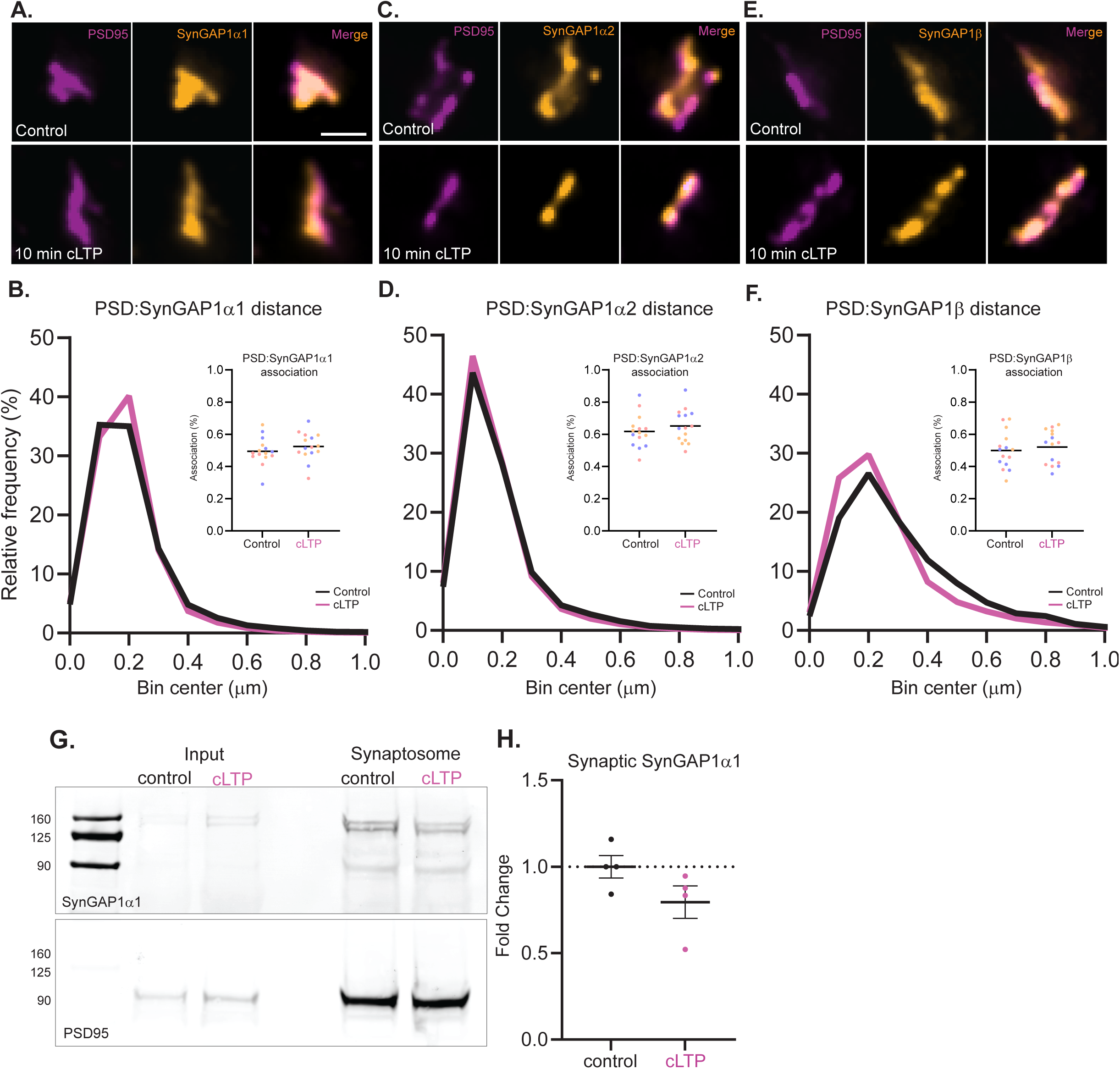
Endogenous SynGAP1 does not disperse at the synaptic level. Representative images and quantification show A) SynGAP1α1 (orange), C) SynGAP1α2 (orange), and E) SynGAPβ (orange) relative to PSD95 (magenta) in control (top panels) and 10 minutes following the induction of cLTP (bottom panels). Graphs show the frequency distribution of **B)** SynGAP1α1, **D)** SynGAP1α2, and **F)** SynGAP1β relative to PSD95. Inset graphs show the proportion of SynGAP1 isoforms colocalising with PSD95. Statistical significance was determined by unpaired t test. Normality for all data was determined by Shapiro-Wilk normality tests. No statistical difference was found between control and cLTP conditions. *n* = 15 FOV, 1840-2729 synapses, 3 biological replicates. Scale bar = 1 µm. **G)** Representative western blot showing SynGAP1α1 (top) and PSD95 (bottom) expression in input (left) and synaptosomal (right) fractions following control and 10 min cLTP conditions. **H)** Quantification of synaptosomal SynGAP1α1 levels following cLTP showed no significant difference to control. *n* = 4 biological replicates. Normality was determined by Shapiro-Wilk normality tests and statistical significance was calculated by Unpaired t test.

### Suppression of SynGAP1α2 protein synthesis following induction of cLTP

Our original hypothesis proposed SynGAP1 synthesis following its dispersion from the synapse as a mechanism to drive SynGAP1 back to the synapse. As we conclusively demonstrate SynGAP1 does not disperse from the synapse, but show evidence that enhanced SynGAP1 expression during LTP increases the risk of excitotoxicity (Supplemental Fig. 3), we sought to assess whether SynGAP1 translation would instead be actively repressed during cLTP. We examined the population of newly synthesised SynGAP1α2 protein 30 minutes (early cLTP) and 60 minutes (late cLTP) following the induction of cLTP (Fig. 6), finding that within the first 30 minutes of cLTP, synthesis of dendritic SynGAP1α2 is significantly reduced compared to controls (Fig. 6 A & C). This effect was found to be both temporary, returning to control levels by 60 minutes, and specific to the dendrites, as somatic newly synthesised SynGAP1α2 remained unchanged compared to controls (Fig. 6 A & B). These results highlight a novel link between synaptic plasticity and translational suppression of negative regulators of plasticity, particularly those that play a key role in tuning synaptic cascades, such as SynGAP1.

**Figure 6.**
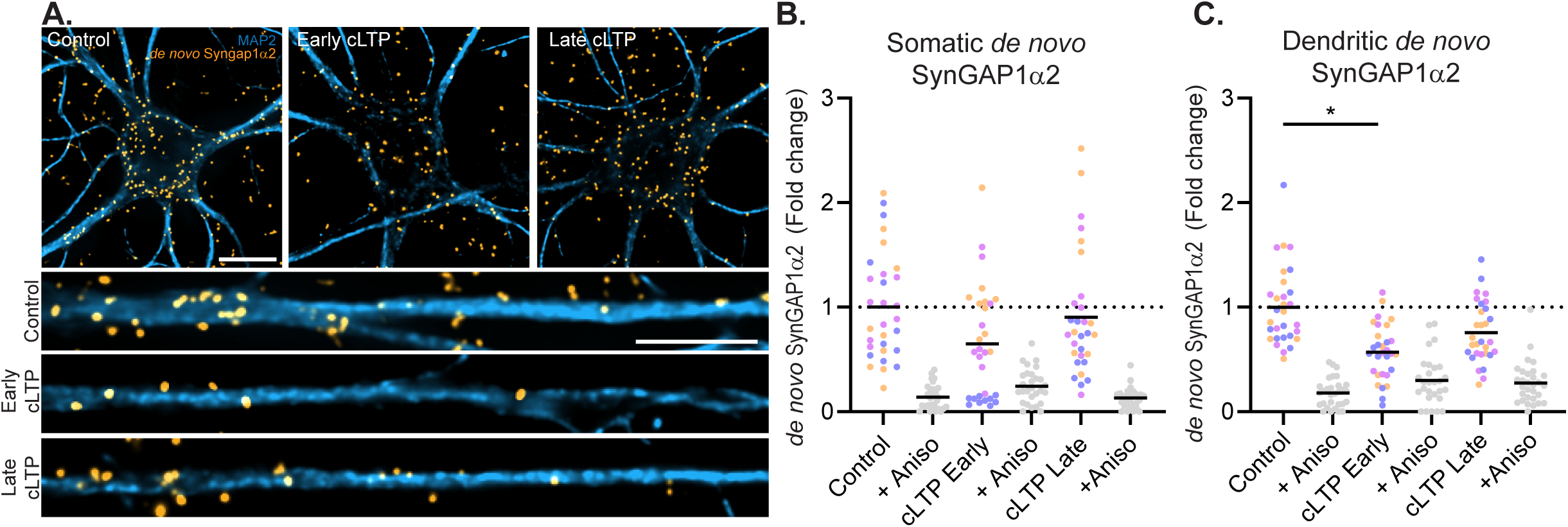
cLTP suppresses SynGAP1α2 synthesis. Representative images show **A)** *de novo* SynGAP1α2 (orange) in the soma (top panels) and dendrites (bottom panels; MAP2, blue) of control, early cLTP and late cLTP conditions. Scale bars = 10 µm. Quantification of **B)** somatic and **C)** dendritic *de novo* SynGAP1α2 expression shows a suppression of dendritic SynGAP1α2 following cLTP. *n* = 28-30 cells, 3 biological replicates. Significance was determined by Ordinary one-way ANOVA or Kruskal–Wallis one-way ANOVA with Dunn’s multiple comparisons test where appropriate. *p* = 0.039*. Normality for all data was determined by Shapiro-Wilk normality tests. All data points are coloured to represent data from 3-4 independent biological replicates.

## Discussion

In this study, we sought to investigate the role of the newly synthesised SynGAP1 protein in the regulation and manifestation of synaptic plasticity. Specifically, we aimed to answer whether local synthesis of SynGAP1 restores the plasticity block alleviated following cLTP-mediated dispersion of SynGAP1 from the synapse. Previous work has shown that immunogold-labelled^30^ and GFP-tagged^20^ SynGAP1 proteins relocalise from the PSD during stimulation. This phenomenon has been described as a necessary component in relieving downstream Ras/Rap inhibition and triggering signalling cascades that induce LTP, including synaptic AMPAR insertion and spine growth.

As an initial step, we attempted to replicate these findings, and explore the timescale for SynGAP1 return to the synapse. While we observed activity-dependent changes in the localisation of GFP-tagged SynGAP1α1, our results revealed a previously unreported level of inconsistency in this response. Rather than a uniform dispersion phenotype, GFP-SynGAP1α1 exhibited a broad spectrum of behaviours across neurons, ranging from no detectable redistribution to widespread, cell-wide dispersion affecting the majority of synapses. This heterogeneity raised questions regarding the robustness and physiological relevance of the previously described dispersion mechanism.

To address this, we examined the behaviour of endogenous SynGAP1 isoforms using both conventional immunofluorescence and super-resolution microscopy. In contrast to the GFP-based observations, we found no significant or quantifiable relocalisation of endogenous SynGAP1 at the PSD following stimulation. While a subtle reduction of SynGAP1 is seen in synaptosomes following cLTP induction, this likely reflects a slight affinity change for the PSD only noticeable when SynGAP1 is removed from its cellular context. These findings suggest that, rather than undergoing large-scale spatial redistribution, endogenous SynGAP1 may instead be regulated through more subtle mechanisms, such as changes in its binding affinity for PSD scaffold proteins like PSD95, or through post-translational modifications that alter its functional state without necessitating physical dispersion.These results are consistent with previous work providing evidence that the key function of SynGAP1 lies in its enzymatic role^41^, and calls into question models that propose SynGAP1 as a structural barrier against AMPAR insertion^41,42^.

Given that endogenous SynGAP1 isoforms did not compare in response to that observed during the GFP-SynGAP1α1 dispersion experiments, it is possible that the range of responses evident may be due to off-target effects of GFP tagging itself. It is widely understood that GFP-tagging of proteins of interest leads to unwanted, largely overlooked artefacts^43–47^, especially for proteins which form condensates^48–50^, including SynGAP1^42^.

We estimate SynGAP1 isoform occupancy at the synapse of approximately 50-60% coverage across three of the primary C-terminal isoforms. While it is currently unknown the degree to which this occupancy overlaps between these three, it is possible that exogenous overexpression of GFP-SynGAP1α1 may drive SynGAP1α1 to mislocalise to synapses previously unoccupied by α1, or occupied primarily by α2 and β. While isoforms α2 and β do not contain the PDZ domain present in SynGAP1α1, all isoforms contain the coil-coil domain crucial for SynGAP1-SynGAP1 trimerisation^51^. The ability of endogenous SynGAP1 isoforms to trimerise with each other, in addition to trimerising with exogenous GFP-SynGAP1α1 may therefore alter the affinity of SynGAP1α1 for PSD and ultimately the propensity of a synapse to disperse. In fact, this trimerisation has been thought to facilitate proper synaptic localisation of SynGAP1^51^. However, we found no clear correlation between the level of synaptic GFP-SynGAP1α1 expression and the degree of dispersion displayed, indicating expression alone may not explain this observation. Together, this may provide rationale for the unreliable dispersion characteristics we observed, and likely represents an artefact of overexpression of a GFP tagged protein^43–47^.

We find the presence of *Syngap1* mRNA at distal dendritic sites, which remained constant throughout dendritic development (Fig. 2). We further provide evidence for the local translation of SynGAP1, confirming previous Ribo-Seq data of the hippocampal neuropil^4^, as well as its regulation by activity, as synthesis of SynGAP1α2, the dominant SynGAP1 isoform, is suppressed during a period of development *in vitro* during which activity is high^36^. A similar suppression of SynGAP1α2 synthesis occurs during early synaptic potentiation, further strengthening the observation of negative translational regulation during periods of elevated synaptic activity. Together, these results identify a previously unseen link between increased synaptic activity and SynGAP1 translational regulation, making SynGAP1 a unique case of translation negatively correlated with activity^52^. We propose the suppression of SynGAP1α2 translation reduces the availability of SynGAP1 during periods of synaptic strengthening, corresponding to periods when pre-existing synaptic SynGAP1 is rapidly inactivated by phosphorylation^20,21^. This translational repression appears to be neuroprotective, as overexpression of SynGAP1 during LTP sensitizes neurons for excitotoxic stress (Supplementary fig. 3), an observation not seen in our SEP-GluA1-expressing control neurons. While GFP-SynGAP1 expression levels were unable to explain this observation alone, it is likely that any level of expression of GFP-SynGAP1, during a period of synaptic activity in which the synthesis of SynGAP1 is typically suppressed, floods the system with excessive levels of SynGAP1 protein, which may decouple the initial Ca^2+^-dependent signaling cascades, such as CaMKII – involved in inactivating SynGAP1 – from the activation of the cascades regulated by SynGAP1 such as Ras. This decoupled state may lead to impaired handling and regulation of Ca^2+^ during periods of heightened activity, leading to excitotoxic stress, which may be consistent with previous reports, showing overexpression of SynGAP1α1 reduces synaptic function^53^, and promotes apoptosis in epithelial cells via suppression of the Wnt/β-catenin pathway ^54^.

Practically, limiting the availability of functional SynGAP1 following the induction of LTP, may help explain the refractory period imposed on potentiated synapses^33^. During this window, synapses are transiently resistant to further potentiation, suggesting the existence of an intrinsic “plasticity brake” that prevents runaway strengthening. This refractory period^33^ – released 45-60 minutes later – aligns with the return of SynGAP1α2 synthesis, suggesting a link between local translational control and synaptic responsiveness. As SynGAP1 acts as a negative regulator of Ras/Rap signaling downstream of NMDA receptor activation, its temporary inactivation following LTP induction would permit sustained signaling necessary for synaptic strengthening. This, coupled with suppression of SynGAP1α2 translation may serve to stabilize this refractory window, ensuring that synapses do not prematurely regain the capacity for additional potentiation. This refractory period is found to be dependent on the availability of PSD95, a core scaffold of the postsynaptic density that anchors signaling complexes and organizes receptor positioning. While our results do not support a direct role for SynGAP1 as a structural barrier to AMPAR insertion, reduced SynGAP1α2 synthesis may indirectly preserve PSD95-dependent structural and signaling states that enforce the plasticity break, allowing proper coordination of the changes in molecular composition required for LTP.

This negative translational regulation of SynGAP1 may also explain why haploinsufficiency of SynGAP1 results in SYNGAP1-related intellectual disability (SYNGAP1-ID). Reduced levels of SynGAP1 is linked to the dysregulation of typical synaptic function, expressing as enlarged, overpotentiated spines^25,19,26^, and is ultimately detrimental to the proper expression of LTP and normal brain development^28^. Activity-dependent translational repression of *Syngap1* mRNA may become maladaptive in SYNGAP1-ID, as it can exacerbate protein insufficiency by establishing a negative feedback loop. Reduced SynGAP1 levels may further restrict the neuron’s capacity to increase SynGAP1 production, thereby preventing compensatory upregulation. Investigation of the molecular mechanisms underlying this negative translational control may identify therapeutic strategies to enhance SynGAP1 synthesis. Targeted disruption of this repression could facilitate activity-dependent restoration of SynGAP1 protein levels, providing a potential approach to restore synaptic function in SYNGAP1-ID.

## Materials and Methods

### Antibodies

| Target | Supplier code | Immuno<br>fluorescence<br>dilution | Fixation<br>conditions | Western blot<br>dilution |
| --- | --- | --- | --- | --- |
| MAP2 | Abcam,<br>#AB92434 | 1:1000-1:2000 | 7 mins methanol<br>-20C or 15<br>minutes 4%PFA<br>in PBS with<br>sucrose |  |
| SynGAP1α1 | CST, (F8I5Q)<br>#34124 | 1:1000 | 7 mins methanol<br>-20°C | 1:1000 |
| SynGAP1α2 | CST, (E4Y6I)<br>#56927 | 1:1000 | 7 mins methanol<br>-20°C | |
| SynGAP1β | CST, (F1D4L)<br>#90217 | 1:1000 | 7 mins methanol<br>-20°C | |
| PSD95 | NeuroMab,<br>#75-028 | 1:1000 | 7 mins methanol<br>-20°C | 1:1000 |
| PSD95 | SySy, #124 011 | 1:1000 | 7 mins methanol<br>-20°C |  |
| PSD95 | SySy, #124 308 | 1:2000 | 7 mins methanol<br>-20°C |  |
| Bassoon | SySy, #141 005 | 1:2000 | 7 mins methanol<br>-20°C |  |
| Biotin | Abcam,<br>#AB201341 | 1:500 | 7 mins methanol<br>-20°C |  |
| Anti Chicken<br>Alexa 488 | Thermo,<br>#A11039 | 1:1000 |  |  |
| Anti Chicken<br>Alexa 568 | Thermo,<br>#A78950 | 1:1000 |  |  |
| Anti Mouse<br>Alexa 488 | Thermo,<br>#A11001 | 1:1000 |  |  |
| Anti Mouse<br>Alexa 568 | Thermo,<br>#A11004 | 1:1000 |  |  |
| Anti Rabbit<br>Alexa 488 | Thermo,<br>#A11008 | 1:1000 |  |  |
| Anti Rabbit<br>Alexa 568 | Thermo,<br>#A11011 |  |  | 1:1000 |
| Anti Guinea Pig<br>Alexa 488 | Thermo,<br>#A11073 |  |  | 1:1000 |
| Anti Guinea Pig<br>Alexa 568 | Thermo,<br>#A11075 | 1:1000 |  |  |
| Anti Guinea Pig<br>Alexa 647 | Thermo,<br>#A21450 | 1:1000 |  |  |
| Anti Guinea Pig<br>CF633 | Cambridge<br>Bioscience,<br>#BT20171-1 | 1:1000 |  |  |
| IRDye 680D | Licor,<br>#926-68071 | 1:1000 |  |  |
| IRDye 800CW | Licor,<br>#926-32210 | 1:1000 |  |  |

### Neuron Prep

The preparation of primary hippocampal and cortical cultures was carried out using a modified version of established protocols^55^. Cortex and hippocampi from P0-P1 Sprague Dawley rat pups (timed pregnant animals ordered from Charles River laboratory) were dissociated using papain (Cambridge Bioscience, #252-10145-1) and plated at a low density on glass-bottomed culture dishes (40,000 or 100,000 cells/dish; Mattek, #P35G-1.5-14-C), and 40,000 cells/coverslip (Scientific Laboratory Supplies, #NPC1613) for immunolabelling experiments, or at 1,000,000 cells/well in a 6-well plate (Sigma, #657160) for biochemical experiments. Cells were cultured in Neurobasal A medium (Life Technologies #10888-022), supplemented with B27 (Life Technologies, #17504-001) and Glutamax (Life Technologies, #35050-061) at 37°C/5% CO_2_ for up to 21 days in vitro (DIV).

### Treatment media

For cLTP and FUNCAT labelling experiments, a basal in-house methionine-, Ca^2+^- and Mg^2+^-free treatment media was made and supplemented with 2-3 mM Ca^2+^, 1mM Mg^2+^ and/or 30mg/L methionine (Formedium, DOC0167) depending on the experimental conditions (outlined in each experimental detail section). The following were added to a 1 L volumetric flask: cell culture-grade H_2_O (Corning, #25-055-CM), adjusted to 260 mOsm and pH 7.4. The media was then sterile-filtered and stored at 4°C or -20°C until use.

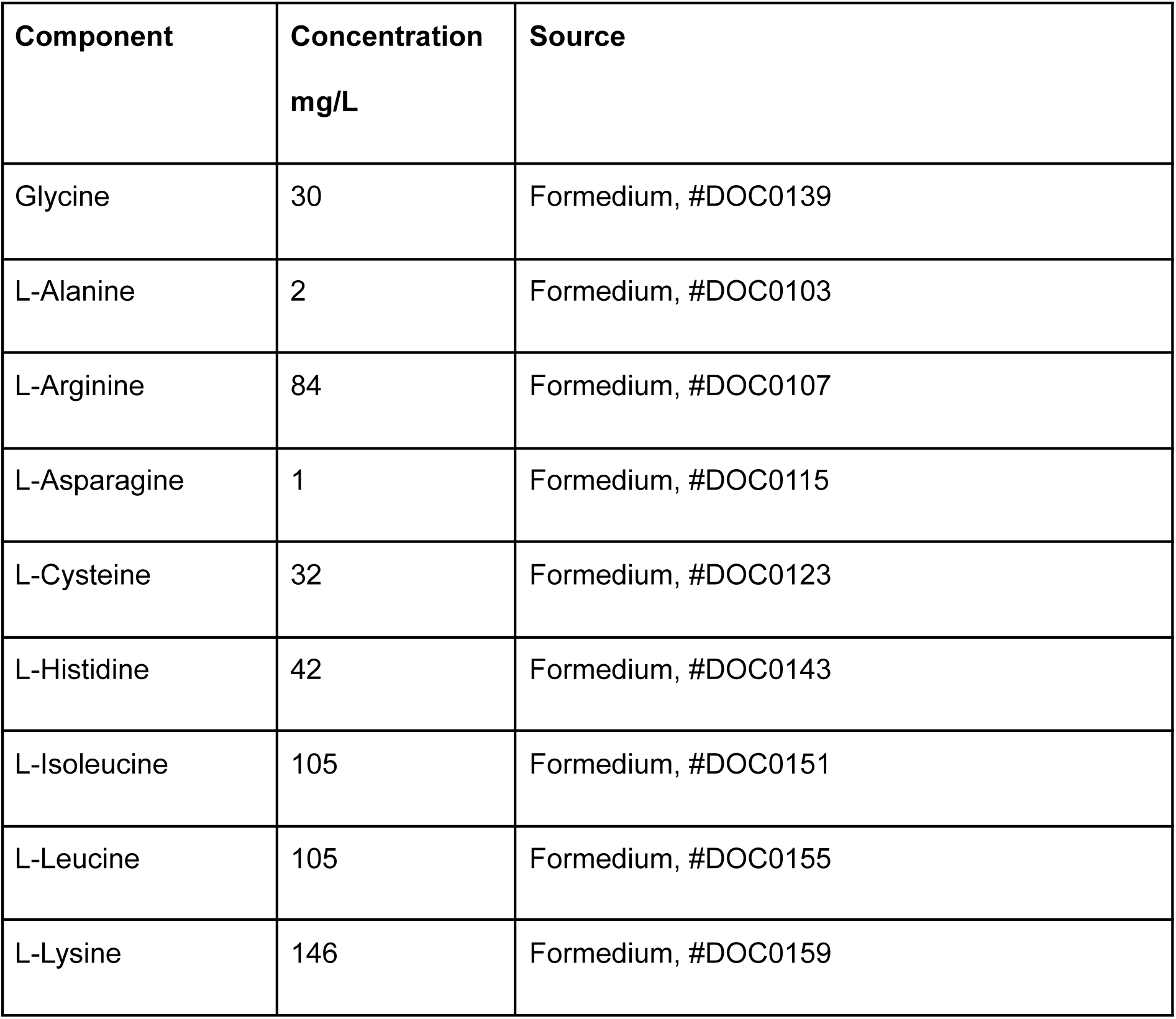
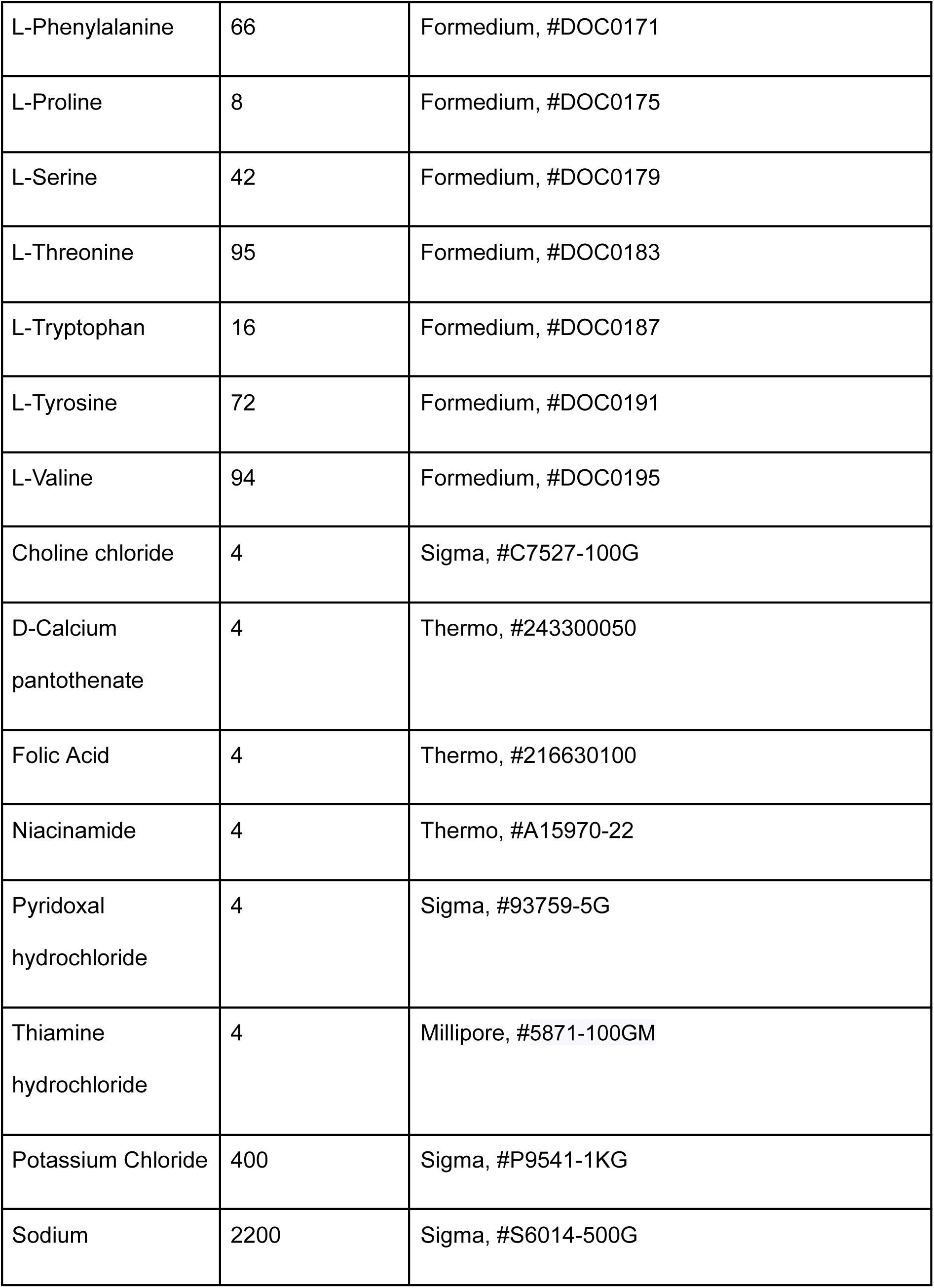
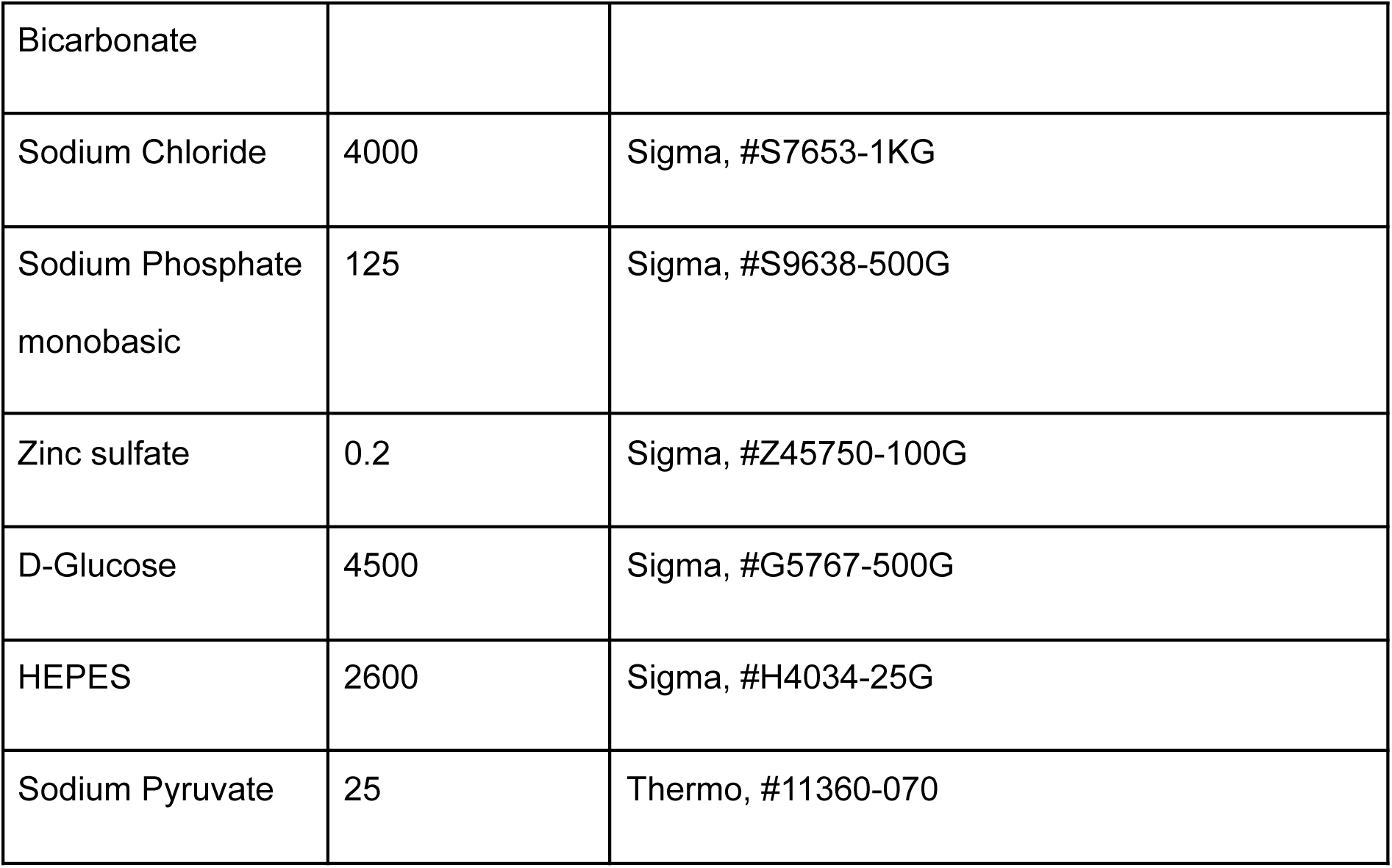

### cLTP

A modified version of the glycine-induced cLTP method^56^ was performed as outlined. Sixteen hours prior to cLTP induction, APV (50 µM; Tocris, #0106) was added to each dish to induce upregulation of N-methyl-D-aspartate receptors^57^. The following day, the media was replaced with homemade treatment media supplemented with B27, Glutamax, 3 mM CaCl_2_, methionine, 200 µM glycine, and 100 µM picrotoxin (Tocris, #1128) for 5 minutes. Media was then removed and replaced with either glia-conditioned Neurobasal A, supplemented with B27 andGlutamax or treatment media supplemented with 2 mM CaCl_2_, 1 mM MgSO_4_, and methionine, and cells were fixed or harvested at the indicated time points post-treatment.

### Immunocytochemistry

To detect proteins in culture throughout development or following cLTP induction, cells were fixed in ice-cold methanol for 7 minutes at -20°C, unless otherwise stated. Samples were permeabilised and blocked in 4% bovine serum albumin (Tocris, #5217) with 0.1% Triton X-100 in phosphate buffered solution (PBS; pH 7.4) for 1 hour at RT. Samples were then probed with appropriate primary antibodies (see table; overnight, 4°C). Following this, the samples were washed 3 x 5 min in PBS (pH 7.4), secondary antibodies were applied for 1 hour at RT, and the samples were then washed 3 x 5 min in PBS (pH 7.4) and mounted for imaging. Cells were imaged using a Nikon Eclipse Ti2 microscope and 60x oil immersion 1.42 NA Olympus objective. Puncta count and intensity analyses were performed using a custom ImageJ script (https://github.com/donlinasplaboratory/Livingstone-RW-2026.git). In brief, 100µm dendritic segments were selected and straightened in ImageJ and the entire soma was outlined. The number and intensity of punctate signals were determined using the ‘Analyze Particles’ plugin.

### RNA FISH

RNA-FISH was performed using the ViewRNA™ ISH Cell Assay Kit (Thermo, #QVC0001) and the manufacturer’s instructions. In brief, cells were fixed in PFA in PBS containing Sucrose (155.42 mM) for 15 min at RT. Cells were then dehydrated in increasing concentrations of EtOH, and stored at -20°C in 100% EtOH until processed for downstream analysis. Following rehydration, samples were permeabilised in Detergent Solution for 5 min at RT, before being washed 2x (Mg^2+^/Ca^2+^ free PBS, pH 7.4), and probed with probe sets for *Syngap1* (VC1-3063080-VCP; 1:100) and *Dlg4 (Psd95;* VC6-10835-VC; 1:100), in Probe Set Diluent at 40°C for 3 hrs. Following incubation, cells were washed in Wash Buffer (3 x 2 min), before application of Pre-Amplifier mix (1:25) at 40°C for 30 min. Following pre-amplification, cells were washed (Wash Buffer, 3 x 2 min), before application of Amplifier mix (1:25) at 40°C for 30 min. Following amplification, cells were washed (Wash Buffer, 3 x 2 min). Finally, Label Probe mix (1:25) was added to the samples and incubated at 40°C for 30 min, before washes (Wash Buffer, 3 x 2 min). For detection of neuronal cell bodies and processes, samples were blocked and incubated with antibodies against MAP2 (overnight at 4°C; see table) and appropriate secondary antibodies (1 hr at RT). Cells were imaged using a Nikon Eclipse Ti2 microscope and 60x oil immersion 1.42 NA Olympus objective. Puncta count analysis was performed using a custom ImageJ script (https://github.com/donlinasplaboratory/Livingstone-RW-2026.git). In brief, dendritic segments were selected and straightened in ImageJ, and the entire soma were outlined. The number of puncta was determined using the ‘Analyze Particles’ plugin.

### FUNCAT-PLA

Labelling of newly synthesised proteins was conducted according to a previously published FUNCAT-PLA protocol^58^. In brief, cells were incubated in 4 mM L-azidohomoalanine (AHA, Cambridge Bioscience #HY-140346A) in the presence or absence of 40 μM anisomycin (Tocris, #1290) following a period of methionine starvation in methionine-free treatment media. For experiments examining *de novo* SynGAP1α2 expression, control samples and samples examining *de novo* SynGAP1α2 through development received 30 minutes of methionine starvation, followed by 30 minutes of AHA incorporation in methionine-free treatment media, before fixation (7 min in methanol at -20°C). For samples examining the effect of cLTP on SynGAP1α2 synthesis, on the day of the experiment, early cLTP samples underwent a 30-minute methionine starvation, followed by a 5-minute cLTP induction (as described above), and were then incubated for 30 minutes in AHA before fixation. Late cLTP samples received 5 minutes of cLTP induction, as above, followed by a 30-minute methionine-starvation period and 30 minutes of AHA incorporation before fixation.

For the detection of *de novo* SynGAP1, azide-labelled newly synthesised proteins were alkylated with biotin-linked alkyne via a copper-mediated click reaction. Click reaction mixture comprised of 200 μM triazole ligand (Tris ((1-benzyl-1H-1,2,3-triazol-4-yl)methyl) amine; TBTA, Sigma #678973), 500 μM TCEP (Tris(2-carboxyethyl)phosphine hydrochloride, Thermo Scientific #10530434), 25 μM Biotin-PEG4-alkyne (Biotin alkyne, 2bScientific, #CCT-TA105) and 200 μM CuSO_4_ (Sigma, 451657) in PBS (pH 7.8), incubated on cells overnight at RT. For the detection of *de novo* SynGAP1, cells were permeabilised and blocked, and incubated overnight at 4°C with anti-biotin and anti-SynGAPα2 antibodies diluted in 4% BSA (Tocris, #5217). Donkey anti-mouse PLA-, and donkey anti-rabbit PLA+ probes were applied, followed by ligation and amplification with Duolink detection reagent Orange (Sigma, #DUO94003) according to the manufacturer’s instructions. Cells were imaged using a Nikon Eclipse Ti2 microscope and 60x oil immersion 1.42 NA Olympus objective. Puncta count analysis was performed using a custom ImageJ script (https://github.com/donlinasplaboratory/Livingstone-RW-2026.git). In brief, dendritic segments were selected and straightened in ImageJ and the entire soma was outlined. The number of puncta was determined using the ‘Analyze Particles’ plugin.

### Transfection

DIV 7 neurons were transfected with GFP-SynGAP1ɑ1^22^ (a gift from Richard Huganir), pCI-SEP-GluR1 (Addgene, #24000), or myr-mScarlet3 (generated in-house, for membrane labelling) using Effectene (Qiagen, #301425), adapting the manufacturer’s recommended protocol as follows. Existing media is removed from the dish (saved and stored in the incubator) and replaced with base neuronal growth media (unconditioned Neurobasal A + B27 + Glutamax), preequilibrated at 37°C overnight. Per dish, .5ug-2ug DNA is mixed with 75uL QC buffer and 4uL enhancer (mixed well by pipetting), and incubated at room temperature (RT) for 10 minutes. Then, 8 µL of Effectene is added (mixed well by pipetting) and incubated at RT for 10 minutes. Then, 85 μL of the mix is added to the dish and returned to the incubator for 1 hour. Finally, all the media and transfection reagents are removed, and the original glia-conditioned media is added back to the neurons. Neurons were maintained until DIV19-22 for downstream imaging.

### Live-cell imaging

Following transfection and maintenance until DIV18-21, APV (50 µM) was added to each dish 16 hours prior to imaging. On the day of imaging, cells were transferred to treatment medium containing APV (50 µM; Tocris, #0106), Ca^2+^, Mg^2+^, and methionine, placed in the imaging chamber (Okolab, #H301-K-FRAME), and equilibrated at 37°C/5% CO_2_. Cells were switched to complete treatment media without APV and were identified by healthy myr-mScarlet3 expression. Following 10 minutes of baseline recording, cells were switched to cLTP induction media (as above) for 5 minutes, after which the media was switched to complete treatment media for 1 hour. Images were collected at 30 second intervals for the duration of each phase of the experiment; each channel was acquired with a 50-ms exposure (2% LED power) across all experiments. To stabilise focus, we used Perfect Focus (Nikon) System on a Nikon Eclipse Ti2 microscope using a 100x 1.45 NA objective. For analysis of GFP-SynGAP1α1 overexpression, baseline fluorescence values were calculated by averaging the final 20 frames of each baseline recording. Excitotoxicity was determined by observation of dendritic fragmentation and membrane swelling (blebbing). To determine single spine intensity of GFP-SynGAP1α1, images were subjected to 2D registration, histogram matching photobleaching correction, and rolling background subtraction. Entire primary dendrites were selected and straightened, and single GFP-SynGAP1α1 puncta were identified with circle ROIs encompassing the entire puncta from the final frame of baseline images. Intensity was measured for each ROI in the final frame of each imaging phase (baseline, induction, recovery).

### Synaptosomes & Western Blot

Isolation of neuronal synaptosomes was performed using Syn-PER™ Synaptic Protein Extraction Reagent (Thermofisher, #87793). In brief, neurons were washed (2x quickly) with PBS (4°C, pH 7.4), Syn-PER™ reagent containing cOmplete™ Protease Inhibitor Cocktail (Sigma, #11697498001) was added to each well, and cells were collected using a cell scraper. The collected material was centrifuged at 1200 × g for 10 minutes at 4°C, and a sample of the homogenate was collected. The remaining supernatant was then centrifuged at 15,000 × g for 20 minutes (4°C). The resulting synaptosomal pellet was resuspended in fresh Syn-PER™ reagent for downstream biochemical analysis. For analysis of whole cell and synaptosomal fractions, aliquots containing 15-20 μg of protein are used for Western blot analysis. Protein extracts were separated by SDS-PAGE (4–12% w/v; Thermo, #NP0321BOX) in MES SDS-PAGE running buffer (Thermo, #NP002) and transferred to nitrocellulose membrane (Thermo, #LC2001) using wet transfer (Thermo, #NP00061). Membranes were probed with appropriate primary and secondary antibodies (see table) and scanned using a LI-COR imaging system. Band intensities representing both SynGAPα1 and PSD95 were quantified using ImageJ, and SynGAPα1 levels were normalised relative to PSD95.

### Expansion microscopy

Tenfold Robust Expansion Microscopy^59^ was adapted for in-gel immunostaining by integrating elements of a published iUExM workflow^60^. **Anchoring**. Samples mounted on coverslips were incubated in an anchoring solution (2% acrylamide [AA], 1.4% formaldehyde [FA] in 1× PBS) at RT for at least 12 hours. **Gelation.** Following anchoring, samples were briefly washed 3-4 times in PBS to remove excess anchoring solution and stored in PBS until preparation of the monomer solution. Prior to gelation, PBS was removed, and the samples were briefly rinsed with distilled water to remove residual salts. Coverslips were blotted to remove excess liquid and floated cell-side down on monomer solution (approximately 50 µL for 5 mm coverslips and 70-100 µL for 12 mm coverslips). Samples were incubated on ice for 30 min, then transferred to a humidified chamber at 37°C for 1 hour to allow polymerisation. Monomer solution consists of 0.1 M sodium acrylate, 2.0 M acrylamide (AA), 50 ppm N, N’-methylenebisacrylamide (bis), PBS (1x), 1.5 ppt APS, and 1.5 ppt TEMED. Sodium acrylate was made as previously described ^59^. **Denaturation.** After gelation, gels were carefully detached from the coverslips by incubating in 2 mL of denaturation buffer (200 mM SDS, 200 mM NaCl, 50 mM Tris base, pH 6.8) in a 6-well plate under gentle agitation. Detached gels were transferred to 15 mL conical tubes containing 5 mL of fresh denaturation buffer and incubated for 1.5 hours at 85°C. **First expansion.** Following denaturation, gels were transferred to double-distilled water (ddH₂O) in Petri dishes. Water was exchanged every 20–30 min until gel expansion reached a plateau. **Blocking and primary antibody staining.** Expanded gels were incubated in blocking buffer for at least 1 hour, then incubated with primary antibodies diluted in blocking buffer overnight at 12°C with gentle shaking. **Washing.** Samples were washed in PBS 4–5 times, with each wash lasting at least 30 min. **Secondary antibody staining**. Secondary antibodies were diluted in blocking buffer and incubated with gels overnight at 12°C with gentle shaking in the dark. **Washing and second expansion.** Gels were washed 4–5 times in PBS for 10 min each, then transferred to ddH₂O for re-expansion. Water was exchanged every 20–30 min until the expansion plateaued. **Imaging.** Fully expanded gels were transferred to poly-D-lysine–coated coverslips in imaging dishes to minimise gel drift. Excess water was gently blotted, and samples were covered with a second coverslip prior to imaging. Gels were imaged using a Nikon Eclipse Ti2 microscope and 60x oil immersion 1.42 NA Olympus objective.

To quantify expansion data, nearest-neighbour distances were calculated using the DiAna “Distance Analysis” plugin in ImageJ/Fiji. Centre-to-centre distances for all objects (not just touching ones) were calculated and used to assess the spatial relationships between the proteins of interest. For expansion-factor quantification, non-expanded control images and expanded images were used to assess Basson-PSD95 centre-to-centre distances. The expansion factor was estimated by normalising both datasets to the average of the non-expanded Basson-PSD95 distance. To assess SynGAP1 association with PSD95, DiAna analysis was performed, assessing only touching objects, and the % PSD95 associated with SynGAP1 was calculated by taking the number of PSD95 puncta touching SynGAP1 over the total number of detected PSD95 per image.

## Contributions

Study design and conception RWL & PGDA Experiments and analysis RWL & PGDA Writing and editing of manuscript RWL & PGDA Supervision of the project PGDA

## Acknowledgments

We thank members of the lab for critical feedback and suggestions. P.G.D.-A. is funded by the Simons Initiative for the Developing Brain (SIDB), a research grant from CureSynGAP1, and a Wellcome Trust Career Development Award.

---

**Supplemental Figure 1.**
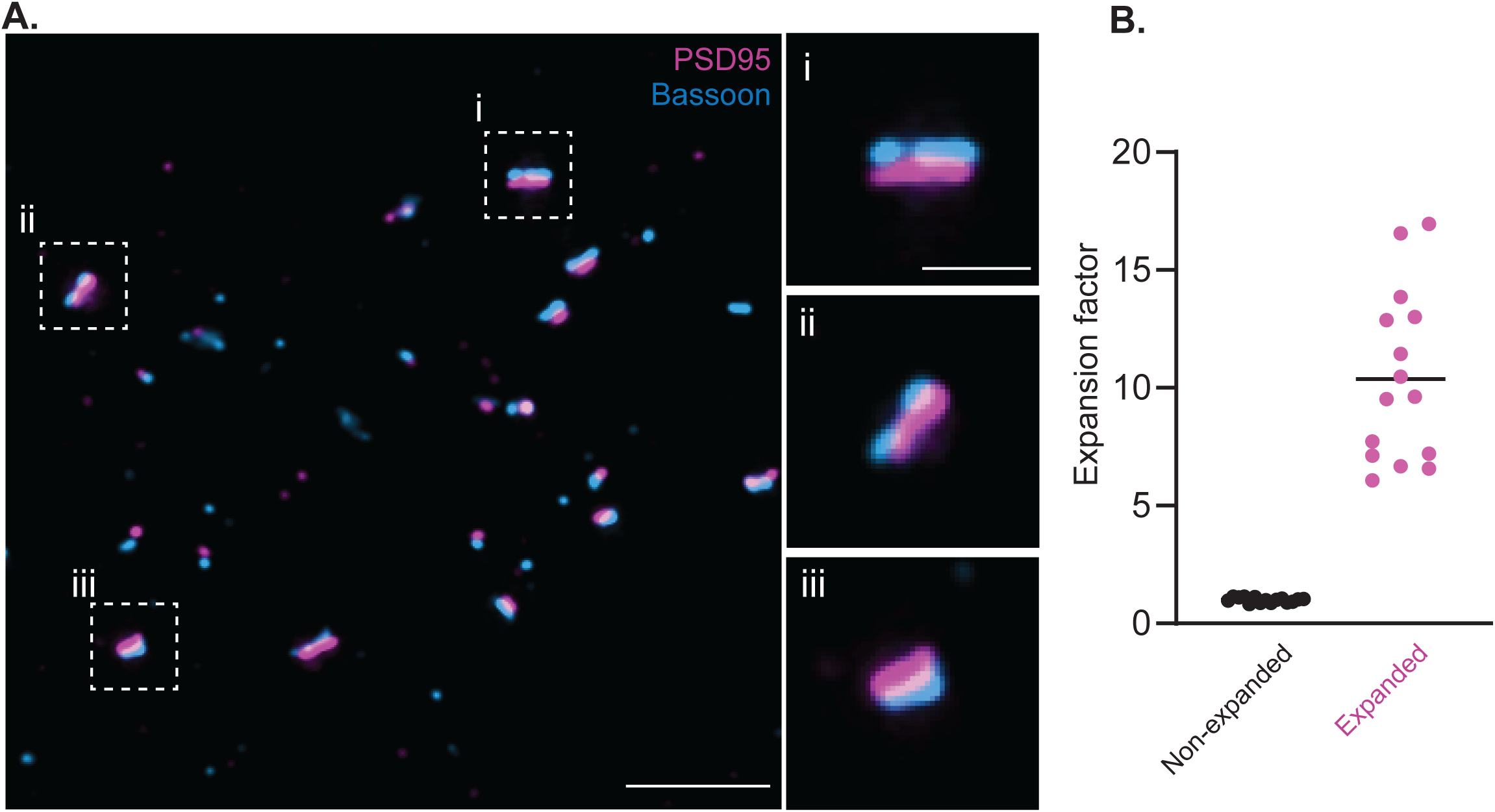
Ten-fold expansion microscopy. Representative images show **A)** a field of view (FOV) of ten-fold expansion of postsynaptic protein PSD95 (magenta) and presynaptic bassoon (blue), scale bar = 5 µm. Insets i-iii show individual examples of ten-fold expanded synapses, scale bar = 1 µm. **B)** Analysis conducted on non-expanded and expanded samples revealed centroid-to-centroid distances of 0.209 ± 0.028 µm and 2.184 ± 0.79 µm for non-expanded versus expanded samples, respectively. This results in an average expansion factor of 10.38 ± 3.62, n = 15 FOV, 3 biological replicates.

**Supplemental Figure 2.**
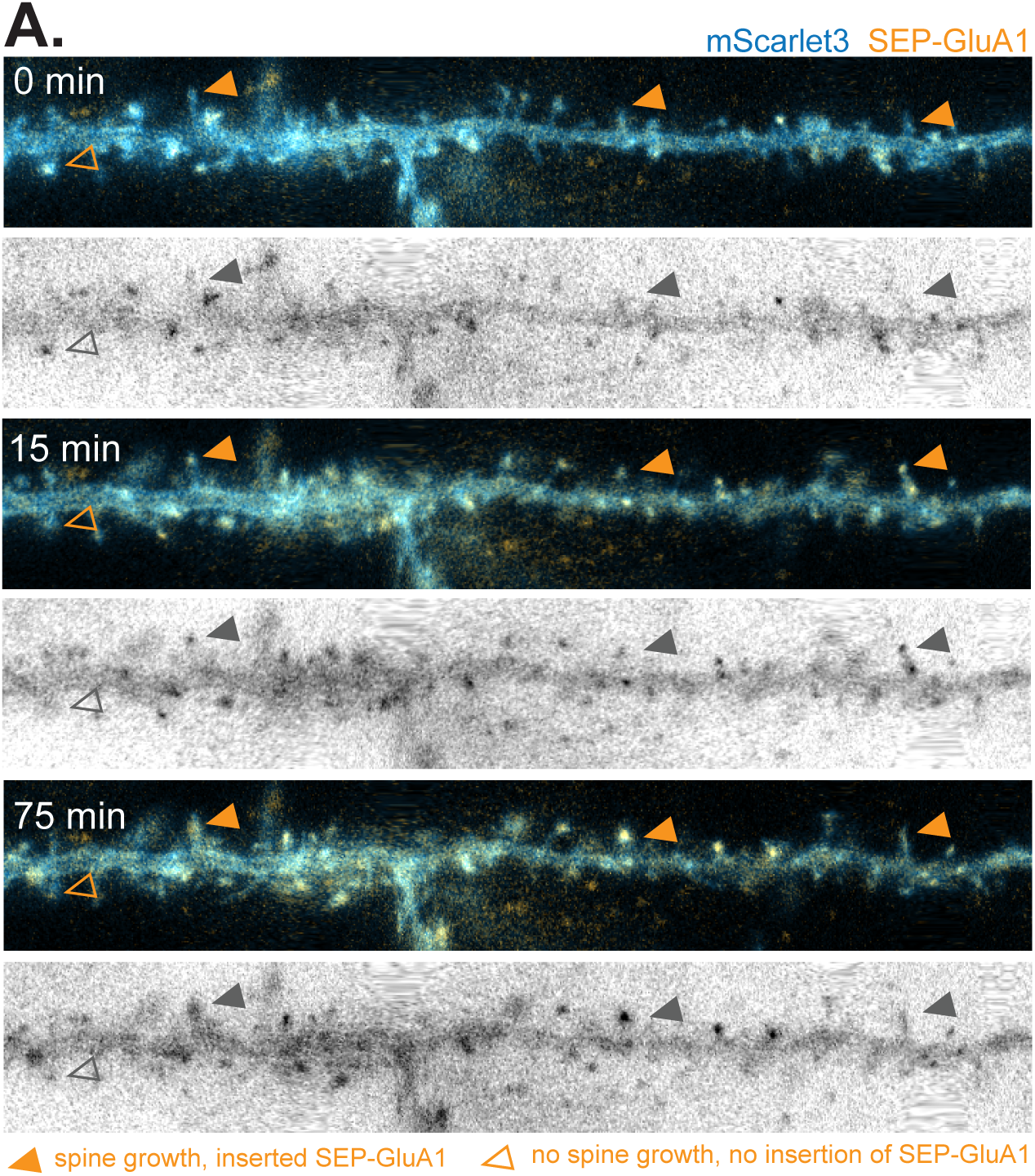
Life cell imaging of SEP-GluA1 confirms the induction of cLTP. Representative time series images of SEP-GluA1 (orange) in mScarlet3-expressing neurons (blue) at baseline (top panels, 0 min), after cLTP induction (centre panels, 15 min), and end of the recovery period (bottom panels, 75 min). Black & white panels show raw SEP-GluA1 signal. Filled orange arrows (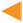) indicate example spines at which GluA1 cell surface exocytosis occurred in the presence of notable spine changes, empty arrows (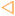) indicate example spines at which GluA1 cell surface exocytosis and spine growth did not occur. *n =* 19 cells.

**Supplemental Figure 3.**
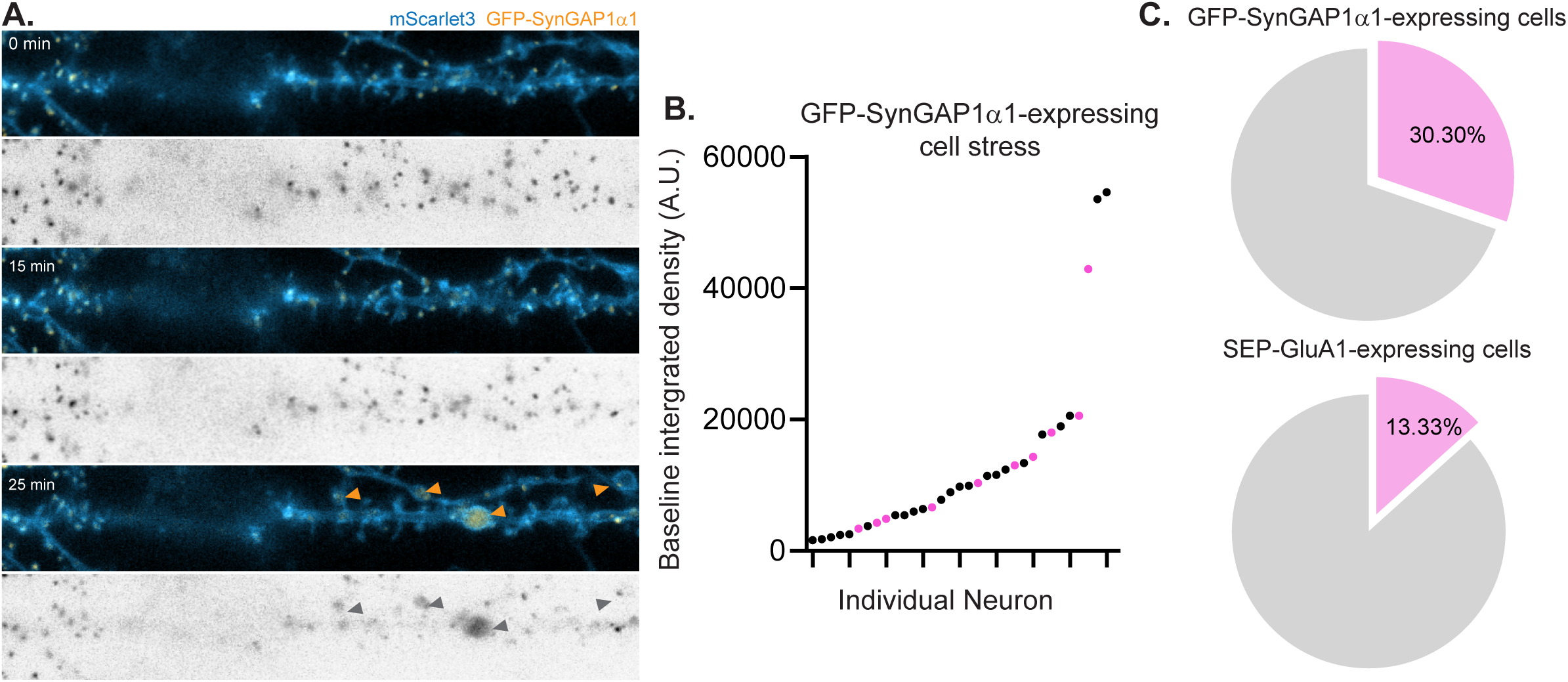
GFP-SynGAP1α1 but not SEP-GluA1 expression leads to increased cell stress during cLTP. **A)** Representative time series images of GFP-SynGAP1α1-expressing (orange) neurons undergoing membrane (blue) beading (blebbing) after the induction of cLTP. Orange arrows (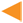) indicate example varicosities following 25 minutes of recording. Black & white panels show raw GFP-SynGAP1α1 signal. **B)** Individual neurons categorized by the observation of cell stress relative to GFP-SynGAP1α1 expression at baseline. *n* = 33 neurons. **C)** Pie charts comparing the proportion of neurons displaying cell stress when expressing GFP-SynGAP1α1 (top) or SEP-GluA1 (bottom).

